# Contraction-dependent intracortical inhibition supports gravity-efficient motor control

**DOI:** 10.1101/2021.06.27.450084

**Authors:** Nicolas Gueugneau, Alain Martin, Jérémie Gaveau, Charalambos Papaxanthis

## Abstract

Efficient control of voluntary movements along the gravity axis requires adapted shifts in muscular contraction modes. In daily life, rising the arm up involves shortening (i.e., concentric) contractions of arm flexors, while the reverse movement can rely on lengthening (i.e., eccentric) contractions of the same muscles with the help of gravity force. Although this muscular-control mode is universal, the neuromuscular mechanisms that subserve the control of such gravity-oriented movements remain unknown. In this study, we designed two neurophysiological experiments that allowed tracking modulations of cortical, spinal, and muscular outputs of arm flexors while healthy humans carried out vertical pointing movements. In conditions where upward and downward movements revealed kinematic features reminiscent of optimal motor commands (i.e., directional asymmetries), we report fine contraction-dependent modulations of the corticospinal output. The overall corticospinal excitability dropped during lengthening contractions (downward movements) compared with shortening contractions (upward movements). Specifically, we did not observe any change in spinal motoneuron responsiveness from cervicomedullary stimulations but a specific increase in intracortical inhibition during lengthening vs. shortening contractions. We discuss these fine contraction-dependent modulations of the supraspinal motor output in the light of feedforward mechanisms that may support gravity-tuned motor control. Generally, these results shed a new perspective on the neural policy that optimize movement control along the gravity axis.

## INTRODUCTION

From sensory perception to movement control, the central nervous system (CNS) has developed efficient strategies to cope with our surrounding *gravity-oriented* environment (White et al. 2020). Whether we stand up or sit on a chair, draw, or reach to grasp an object, the trajectories of unconstrained movements systematically show directional asymmetries. Precisely, upward movements have a shorter time to peak velocity, greater peak acceleration, and larger path curvature than downward movements (Gaveau and Papaxanthis, 2011; Gaveau et al., 2014, 2021; Papaxanthis et al. 1998, 2003, 2005; Yamamoto and Kushiro 2014). A previous study in the journal has demonstrated that this behavior is optimal to save muscle effort (Gaveau et al. 2016). More, it suggests that such efficient control emerge from brain processes.

The shift in muscle contraction modalities along the vertical axis is also a basic feature of movement control in the gravity field. Indeed, naturally paced upward and downward arm movements required specific activations of the flexors/antigravity muscles, i.e., *shortening-concentric* contraction when moving upwards (against gravity) and *lengthening-eccentric* contraction when moving downwards (with gravity). Such a control scheme is not possible in the horizontal plane, where the activation of both agonist and antagonist muscles is necessary to move leftwards or rightwards (Papaxanthis et al., 2003; Gaveau et al. 2021, Hooper, 2012). We recently scrutinized muscular activation patterns during single-degree-of-freedom arm movements in various directions. Using a well-known decomposition method of tonic (muscle activity needed to keep the arm still against gravity) and phasic (muscle activity needed to accelerate and decelerate the arm) electromyographic (EMG) activities, we demonstrated that phasic electromyograms present systematic negative phases (i.e., phases where EMG activity is less than necessary to compensate gravity torque). This negativity reveals an optimal motor plan where the motor system harvests the mechanical effects of gravity to accelerate downward and decelerate upward movements (Berret et al., 2008; Gaveau et al. 2021).

Previous studies modeled these robust kinematic and EMG patterns as an optimal motor planning policy by which the central nervous system purposely takes advantage of the gravity force to save muscle effort (Berret et al. 2008, 2011; Crevecoeur et al. 2009; Gaveau et al. 2014, 2016, 2021). Such models, deriving from neurophysiological knowledge obtained in human and non-human primates, assume that the brain computes a gravity internal model (Angelaki et al., 2004; Laurens et al. 2013, 2016; Indovina et al. 2005). This brain internal model is then supposed to support optimal motor planning and control. However, despite the abundant behavioral evidence for optimal control of gravity-oriented movements, its neural implementation remains unknown. The contribution of supraspinal and spinal neural mechanisms subserving the shortening and lengthening muscle patterns in the gravity field is unknown. In *constrained* motor tasks, such as maximum force generation, shortening and lengthening muscle contractions result from specific neuromuscular control strategies (Duchateau & Enoka, 2016). Precisely, it has been shown that spinal mechanisms contribute to partially inhibit the neural drive originating from the motor cortex (measured by the size of motor evoked potential, MEP) during lengthening compared with shortening contractions (Duclay et al. 2011, 2014; Gruber et al., 2009, Abbruzzese et al., 1994; Sekiguchi et al., 2001). Gruber et al. (2009) demonstrated a greater reduction of CMEP amplitude (cervicomedullary motor evoked potential) than MEP amplitude during lengthening contractions of the elbow flexor muscles. As CMEP does not involve cortical neurons, while MEP includes both cortical and spinal neurons, it provides a direct assessment of spinal cord motoneurons’ responsiveness to synaptic inputs (Nielsen and Petersen, 1994; Taylor, 2006). Yet, in *unconstrained* motor tasks, the neural drive of shortening and lengthening muscle contractions has not been investigated. It is known that the control of arm dynamics involves both spinal and supraspinal mechanisms. For example, Gritsenko et coll. (2011) found that transcranial magnetic stimulation (TMS) responses in shoulder and elbow muscles changed when interaction torques were resistive but not assistive to arm movements, suggesting that the descending motor command includes compensation for passive limb dynamics. Kurtzer et al. (2008), during perturbed arm movements, found that short-latency reflexes of shoulder muscles were exclusively linked to shoulder motion, whereas long-latency reflexes were sensitive to both shoulder and elbow motion. They concluded that long-latency reflexes possess an internal model of limb dynamics, a degree of motor sophistication that was previously reserved to voluntary motor control. For postural control of the upper limb, recent work shows that even short latency feedback loops (20ms, i.e. eminently spinal) are tuned to the motor context, thereby producing sophisticated efficient motor control (Weiler et al. 2019, 2021).

In this study, we investigated whether supraspinal and/or spinal mechanisms contribute to the kinematic and EMG features of arm movements performed along the vertical axis. To do so, we conducted two experiments where participants performed upward and downward pointing movements. Non-invasive neurophysiological methods were used to finely track the modulations of cortical, spinal, and muscular outputs of arm flexors during the lengthening and shortening contraction phases of arm motion along the vertical axis.

## RESULTS

### Arm kinematics

Sixteen healthy participants performed single-joint, visually guided pointing movements in the parasagittal plane with their right forearm (rotation around the elbow; see Figure 1, left panels). We used one degree of freedom movements to isolate the mechanical effects of gravity (Gentili et al. 2007; Le Seac’h and McIntyre 2007; Gaveau et al. 2014). Precisely, during single-joint vertical forearm movements, inertia (i.e., the distribution of the forearm mass around the elbow joint in a body-fixed coordinate system) remains constant, and inertial torque is only related to joint acceleration. All participants accomplished downward and upward movements at a comfortable speed by mobilizing the elbow joint (no rotation at the other joints) without deviating from the sagittal plane (see Methods-Data analysis). Motion capture techniques were used to track movement kinematics and revealed that velocity profiles were single-peaked and bell-shaped (Fig 1, right panels).

**Figure 1.**
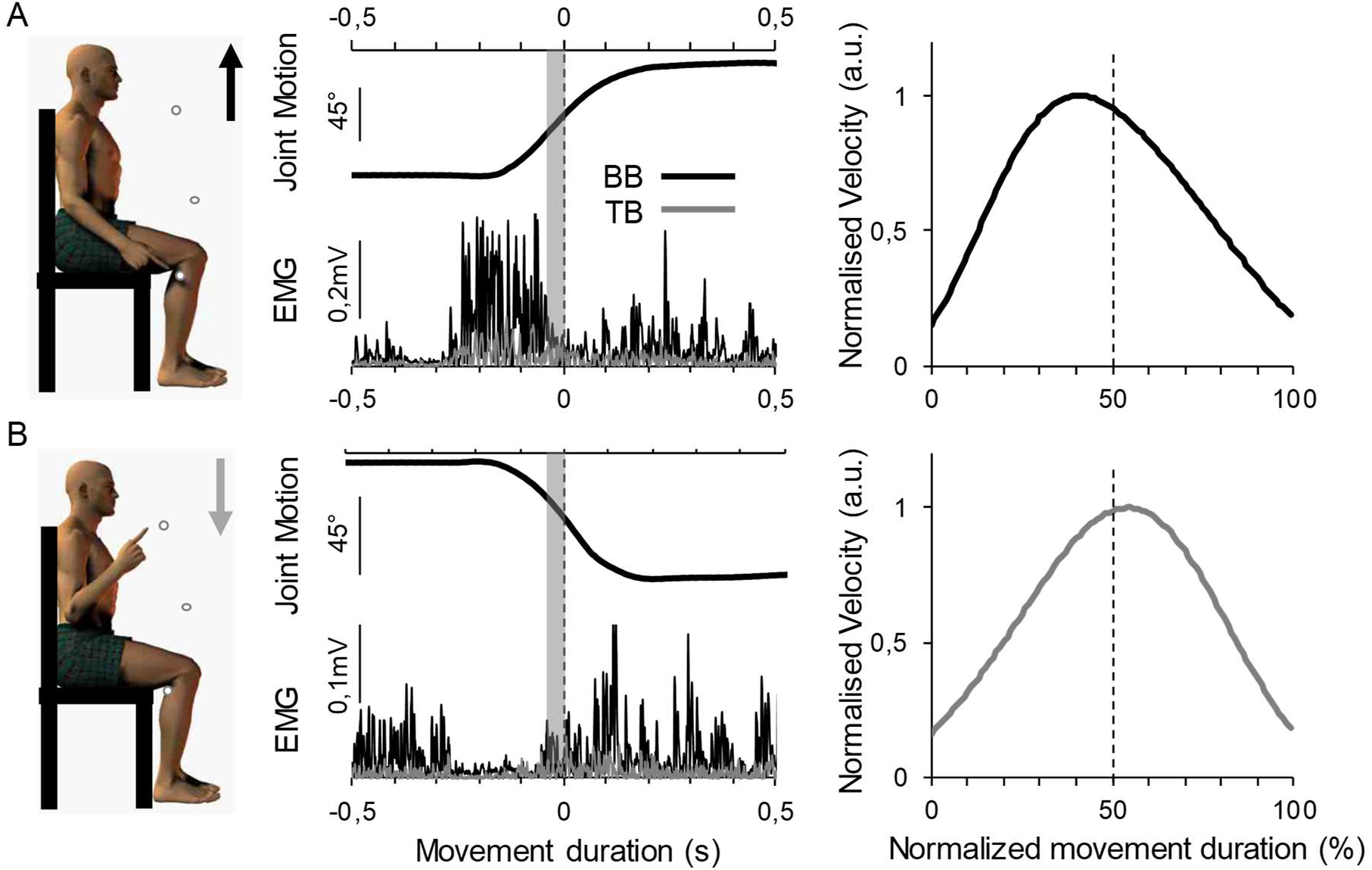
Kinematic and EMG determinants of the motor task. **A.** Schematic representation of a participant executing an upward movement (left panel), with the corresponding temporal evolution of joint motion and EMG activity during the movement (middle panel). Joint motion is represented as the upper arm-forearm angle, and EMG activity is represented as the rectified EMG signal for the BB (black trace) and the TB (grey trace). Typical normalized velocity profile (right panel). **B.** Same as A, but for a downward movement. For the middle panels, the vertical dotted lines show the time of electrophysiological stimulations, and the shaded grey area indicates the time window during which the pre-stimulus EMG activity was quantified (see *Data analysis*). For the right panel, the vertical dotted line indicates the mid-movement time and allows to appreciate the directional asymmetry, with a clear shift of peak velocity toward the beginning and the end of the motion, for upward and downward movements, respectively.

Average values 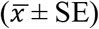 of the main forearm kinematics are shown in Table 1. Movement amplitude, duration, and mean velocity did not show any significant difference between upward and downward movements, while peak velocity and acceleration were higher (acceleration was close to significance; *P* = 0.050) during upward compared to downward movements. Also, we observed significant differences for the relative time to peak acceleration (rTPA) and the relative time to peak velocity (rTPV), with lower values for upward movement compared to downward movements. Overall, these specific directional asymmetries confirm previous results from the literature that were shown to reveal an optimal motor control process minimizing muscle effort in the vertical plane (Berret et al. 2008, 2011; Gaveau et al. 2014, 2016, 2021).

**Table 1.** Effect of movement direction on forearm movement kinematics, and direction ratios (i.e. invariant parameters). rTPV = relative time to peak velocity; rTPA = relative time to peak acceleration.

|  | Movement |  | *P-value* | Cohen's *d* |
| --- | --- | --- | --- | --- |
|  | *Type of muscle contraction* |  |  |  |
|  | Downward<br>*Lengthening* | Upward<br>*Shortening* |  |  |
| *Kinematic features* |  |  |  |  |
| Amplitude (m) | 0.52 ± 0.11 | 0.51 ± 0.12 | 0.178 | 0.08 |
| Duration (s) | 0.5 ± 0.03 | 0.49 ± 0.03 | 0.972 | < 0.001 |
| Mean Velocity (m/s) | 1.16 ± 0.09 | 1.15 ± 0.07 | 0.963 | 0.12 |
| Peak Velocity (m/s) | 1.72 ± 0.11 | 1.83 ± 0.15 | 0.040 | 0.83 |
| Peak Acceleration (m/s<sup>2</sup>) | 9.25 ± 1.26 | 9.76 ± 1.19 | 0.050 | 0.41 |
| *Movement direction ratios* |  |  |  |  |
| rTPV | 0.54 ± 0.04 | 0.49 ± 0.04 | 0.035 | 1.11 |
| rTPA | 0.21 ± 0.02 | 0.17 ± 0.01 | 0.036 | 0.97 |

Data are mean ± SE

### Neurophysiological parameters - Experiment A

Corticospinal excitability was evaluated by using single-pulse TMS, short intracortical inhibition (SICI) by paired-pulse TMS, and silent period (SP). Spinal excitability was evaluated by cervicomedullary stimulation (CMS), while muscle excitability was assessed through Mmax recordings. The stimulations were applied when the elbow joint reached 90° during upward or downward movements, which allowed us to precisely target the specific contraction modalities of the BB.

At the muscular level, the motor task allowed us to focus on the BB which was activated in the two contraction modalities according to movement direction; i.e., lengthening and shortening contractions for downward and upward movements, respectively. Electromyographic signals (EMG) from both the BB and the Triceps Brachii (TB) of the right arm were recorded during the whole movement course. Fig 1 (middle panels) qualitatively illustrates the activation patterns of both muscles during the motor task. It could be noticed that upward and downward forearm movements are mainly produced by the activation of the BB. BB and TB muscle activity patterns were also characterized by computing the root mean square (RMS) of the EMG signals.

Average values 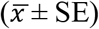 for the main neurophysiological parameters during downward and upward movements are depicted in Table 2 (upper part). M_max_, RMS/M_max_, and muscle coactivation were not significantly different between the two movement directions (in all, *P* > 0.05). This indicates that at an elbow angle of 90° the muscular activity was comparable between movement directions and confirms that forearm motion was mainly controlled by the BB during both lengthening (downward movement) and shortening (upward movement) contractions (as revealed by similar values of coactivation). Notably, this implies that further spinal and supraspinal modulations could not result from differences in muscle activation state between the two movement directions.

**Table 2.**
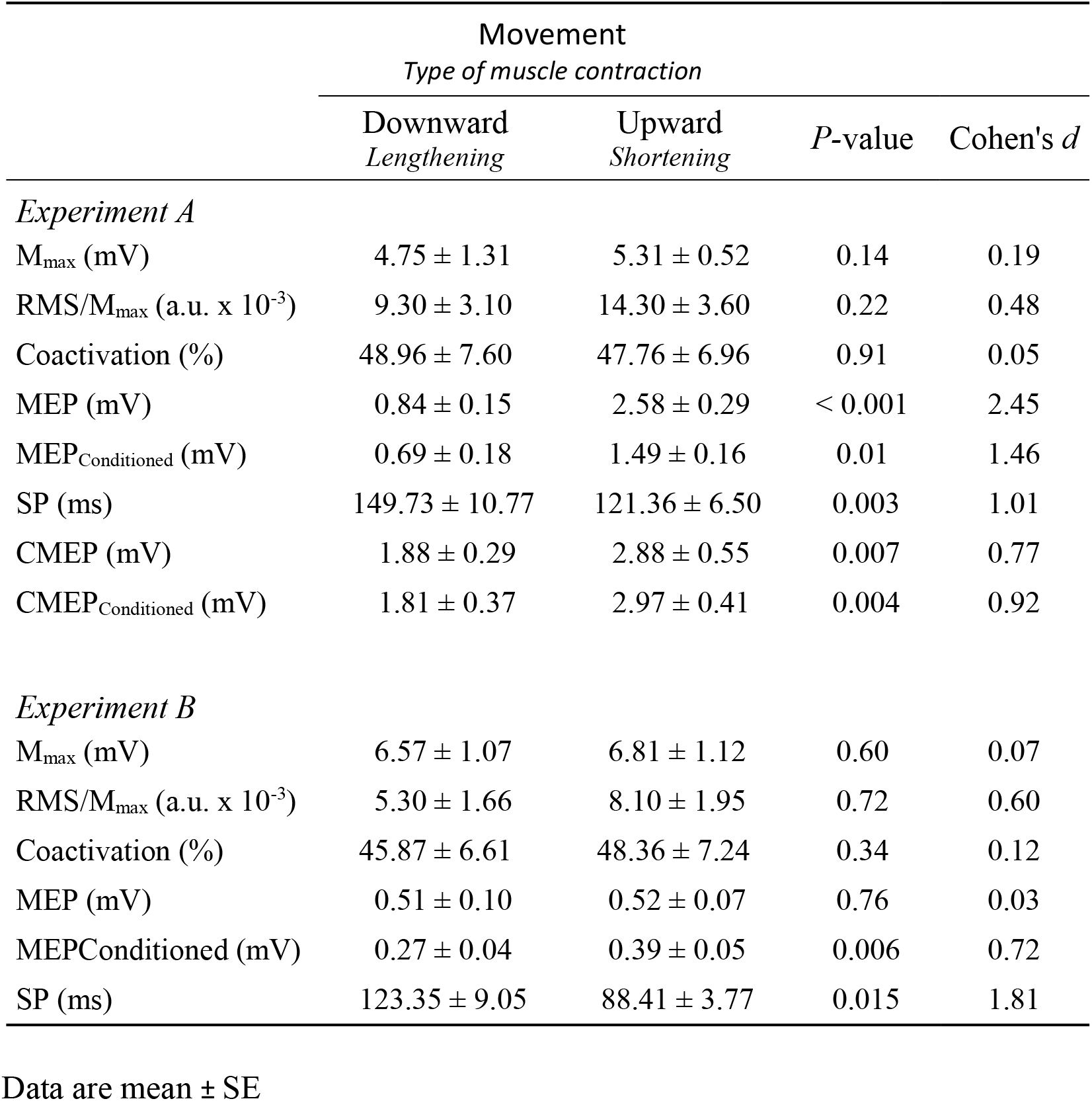
Effect of movement direction on the neurophysiological parameters for experiment **A** (upper part) and **B** (lower part).

|  | Movement |  | *P*-value | Cohen's *d* |
| --- | --- | --- | --- | --- |
|  | *Type of muscle contraction* |  |  |  |
|  | Downward<br>*Lengthening* | Upward<br>*Shortening* |  |  |
| *Experiment A* |  |  |  |  |
| M<sub>max</sub> (mV) | 4.75 ± 1.31 | 5.31 ± 0.52 | 0.14 | 0.19 |
| RMS/M<sub>max</sub> (a.u. x 10<sup>-3</sup>) | 9.30 ± 3.10 | 14.30 ± 3.60 | 0.22 | 0.48 |
| Coactivation (%) | 48.96 ± 7.60 | 47.76 ± 6.96 | 0.91 | 0.05 |
| MEP (mV) | 0.84 ± 0.15 | 2.58 ± 0.29 | < 0.001 | 2.45 |
| MEP<sub>Conditioned</sub> (mV) | 0.69 ± 0.18 | 1.49 ± 0.16 | 0.01 | 1.46 |
| SP (ms) | 149.73 ± 10.77 | 121.36 ± 6.50 | 0.003 | 1.01 |
| CMEP (mV) | 1.88 ± 0.29 | 2.88 ± 0.55 | 0.007 | 0.77 |
| CMEP<sub>Conditioned</sub> (mV) | 1.81 ± 0.37 | 2.97 ± 0.41 | 0.004 | 0.92 |
| *Experiment B* |  |  |  |  |
| M<sub>max</sub> (mV) | 6.57 ± 1.07 | 6.81 ± 1.12 | 0.60 | 0.07 |
| RMS/M<sub>max</sub> (a.u. x 10<sup>-3</sup>) | 5.30 ± 1.66 | 8.10 ± 1.95 | 0.72 | 0.60 |
| Coactivation (%) | 45.87 ± 6.61 | 48.36 ± 7.24 | 0.34 | 0.12 |
| MEP (mV) | 0.51 ± 0.10 | 0.52 ± 0.07 | 0.76 | 0.03 |
| MEP<sub>Conditioned</sub> (mV) | 0.27 ± 0.04 | 0.39 ± 0.05 | 0.006 | 0.72 |
| SP (ms) | 123.35 ± 9.05 | 88.41 ± 3.77 | 0.015 | 1.81 |

Data are mean ± SE

Interestingly, both MEP and CMEP amplitudes were significantly lower for downward vs. upward movements (for both, *P* < 0.05). Figure 2A shows typical MEP and CMEP for downward and upward contractions, while Figure 2B shows normalized MEP and CMEP amplitudes for the same conditions. Though normalized MEP amplitudes were significantly reduced for downward compared to upward movements (*P* = 0.001; *d* = 1.07), no significant difference was found for CMEP amplitudes (*P* = 0.89; *d* = 0.12). The SP was significantly shorter during upward movements compared to downward movements. Because the SP duration can be influenced by the size of the MEP, we normalized the SP duration values by computing the ratio SP/MEP. Again, the normalized SP duration (Fig 2C) was significantly shorter during upward compared to downward movements (*P* < 0.001; *d* = 2.46). Finally, SICI was more pronounced during upward compared to downward movements (Fig 2D; respectively −22.85 ± 9.25 % and −42.45 ± 8.64 % for downward and upward movements; *P* = 0.04; *d* = 0.69).

**Figure 2.**
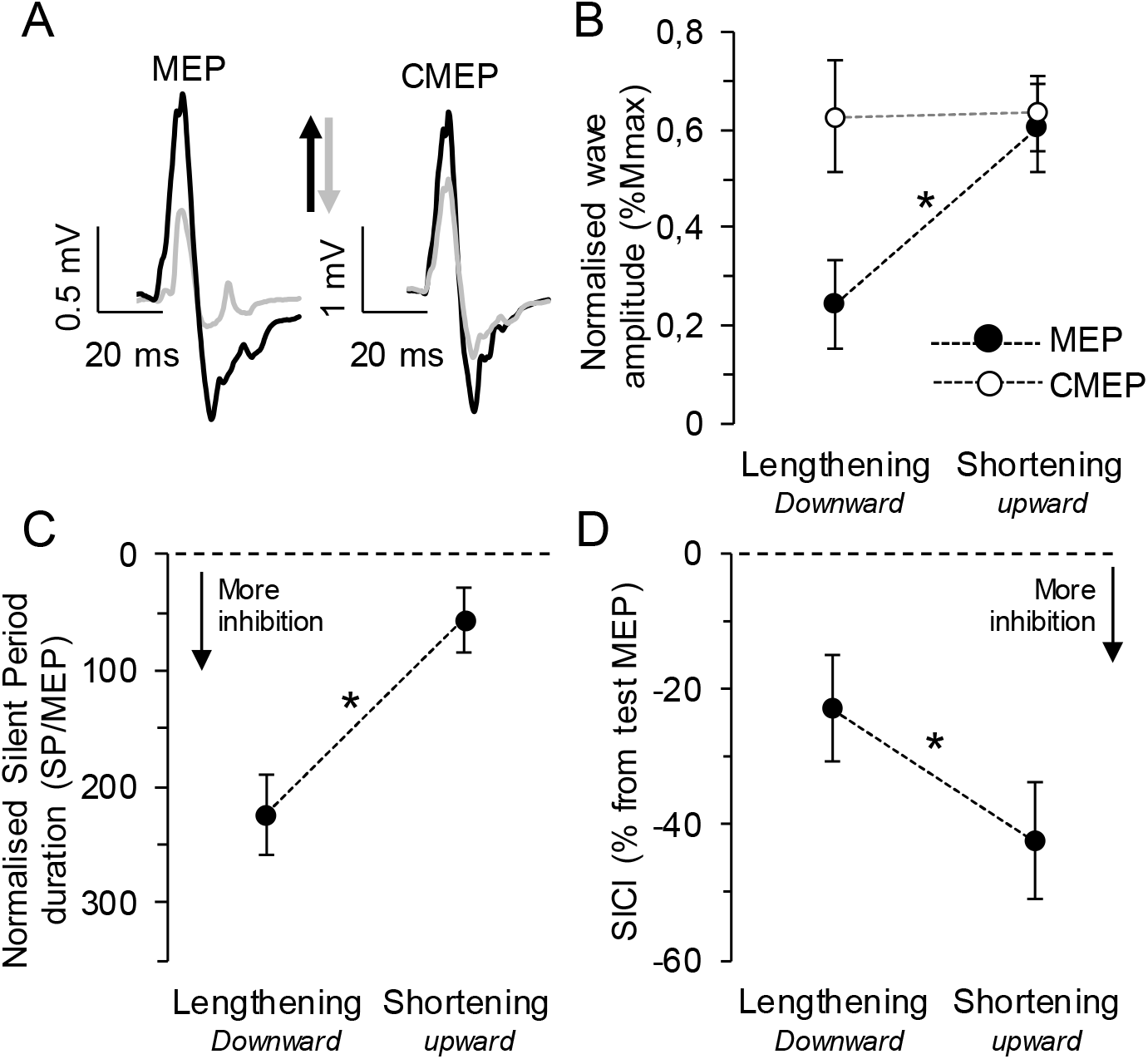
Changes in corticospinal responses and intracortical inhibition during downward (lengthening contraction) and upward (shortening contraction) movements. **A.** Typical MEP and CMEP for downward and upward movements (grey and black traces, respectively). **B.** Mean normalized MEP amplitudes (± SE) of MEP and CMEP (full and empty circles, respectively) for downward and upward movements **C.** Normalized SP duration (± SE) for downward and upward movements. **D.** Mean SICI values (± SE) for downward and upward movements. * Significant difference at *P* < 0.05.

Figure 3A shows typical CMEP_unconditioned_ and CMEP_conditioned_ signals during both downward and upward movements and Figure 3B shows normalized CMEP_conditioned_ amplitudes for the same conditions. Normalized CMEP_conditioned_ amplitudes were not significantly different between contractions (*P* = 0.83; *d* = 0.09). These data come from a paired-pulse TMS-CMS protocol that was specifically designed to evaluate the potential influence of the conditioning (sub-threshold) TMS pulse on spinal motoneuronal excitability (see Methods).

**Figure 3.**
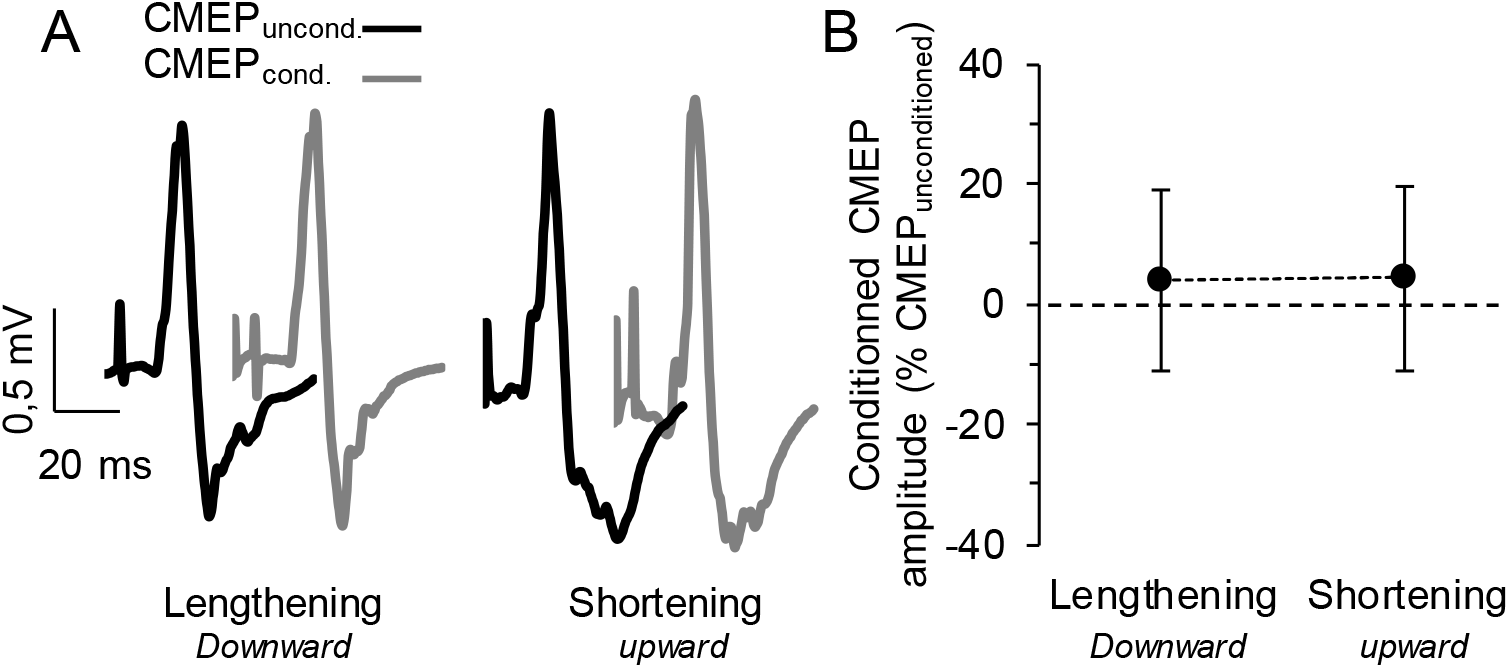
Effect of a conditioning TMS-pulse on CMEP response (paired-pulse TMS-CMS protocol). **A.** Typical unconditioned and conditioned CMEP (black and grey traces, respectively) during downward (lengthening contraction) and upward (shortening contraction) movements. **B.** Mean normalized conditioned CMEP amplitudes (± SE) for both movement directions.

In summary, Experiment A showed a decrease in corticospinal excitability during lengthening (downward movement) vs. shortening (upward movement) contractions, as evidenced by clear modulations of MEP amplitudes; while spinal motor neuron excitability remained similar during both contraction types (no CMEP variation when normalized to M_max_). However, intracortical inhibition showed paradoxical results whether it was assessed by SP or paired-pulse protocol (SICI). It increased during lengthening vs. shortening contractions as measured by SP duration, while it showed the opposite pattern when measured by SICI.

### Neurophysiological parameters - Experiment B

Because the size of the test MEP, in SICI protocols, can bias the magnitude of intracortical inhibition (Sanger et al., 2001; Daskalakis et al., 2002), normalization methods are to be employed to control this size-effect and better characterize SICI mechanisms in motor control tasks (Lackmy and Marchand-Pauvert, 2010; Opie and Semmler, 2014). Accordingly, experiment B was designed to assess intracortical inhibition while controlling corticospinal excitability across the experimental conditions, where test MEPs in the SICI protocol were carefully matched between shortening and lengthening contractions.

Average 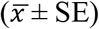 values for the main neurophysiological measures during downward and upward movements are given in Table 2 (lower part). As in experiment A, M_max_, RMS/M_max_, and muscle coactivation were not significantly different between movement directions (in all, *P* > 0.05), further confirming a comparable muscular activity during both movement directions, and a similar agonist/antagonist strategy (as revealed by similar values of coactivation). No significant difference was found for MEP amplitudes during downward and upward movements, indicating an appropriate methodology for matching corticospinal excitability between conditions; yet, MEP_conditioned_ amplitudes were significantly lower during downward compared to upward movements (see Table 2).

Figure 4A shows typical MEP during downward and upward movements, while figure 4B shows normalized MEP amplitudes. The latter were not significantly different between movement directions (*P* > 0.05; *d* = 0.08). The SP was significantly shorter during upward vs downward movements (*P* = 0.015; *d* = 1.81). Also, the normalized SP duration (Fig 4C) was significantly shorter during upward compared to downward movements (*P* = 0.03; *d* = 0.73). SICI showed a similar pattern, therefore opposite to what was observed in Experiment A, as it was more pronounced during downward compared to upward movements (Fig 4D). The inhibition was −24.35 ± 4.68 % and −45.01 ± 5.12 % for upward and downward movements respectively (*P* = 0.03; *d* = 1.48).

**Figure 4.**
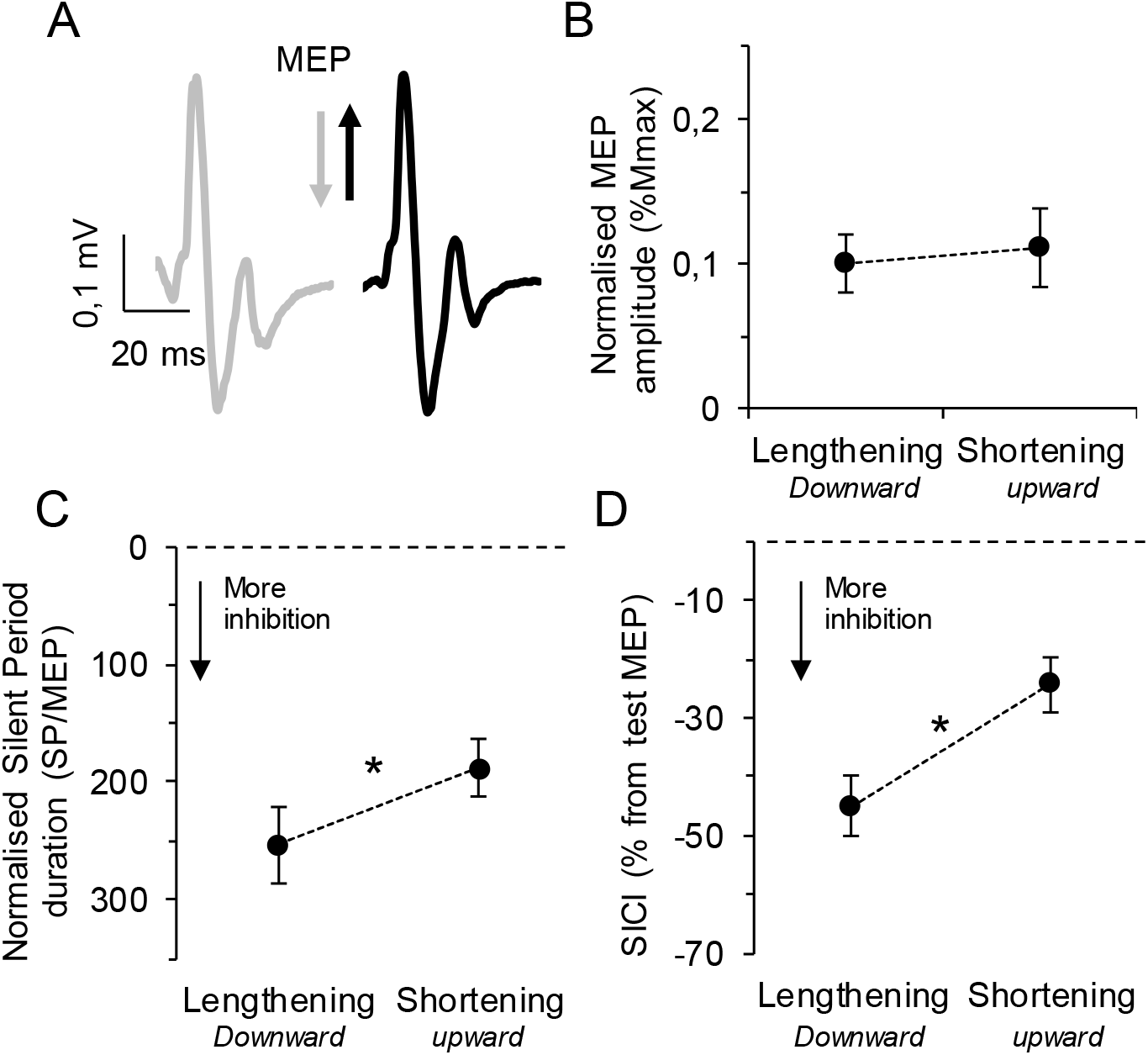
Changes in intracortical inhibition while controlling the corticospinal output. **A.** Typical MEP for downward and upward movements (grey and black traces, respectively). **B.** Mean normalized MEP amplitudes (± SE) for downward and upward movements. **C.** Normalized SP duration (± SE) for downward and upward movements. **D.** Mean SICI values (± SE) for downward and upward movements. * Significant difference at *P* < 0.05.

To summarize, when controlling for test MEP amplitude between directions, Experiment B revealed similar patterns of intracortical inhibition whether it was assessed by SP or paired-pulse protocol (SICI). Precisely, intracortical inhibition was more pronounced during downward (lengthening contractions) than upward movements (shortening contractions).

## DISCUSSION

We investigated the neural mechanisms underlying the control of unconstrained gravity-oriented forearm movements. We designed two experiments that allowed tracking the modulations of cortical, spinal, and muscular outputs of arm flexors while healthy adults accomplished movements along the vertical axis. We confirmed, in line with previous studies, that velocity profiles revealed consistent direction-dependent asymmetries (upward vs. downward). Additionally, forearm movements were performed by activating the flexor muscles only: downward movements (with gravity) were generated by lengthening (eccentric) contraction while upward movements (against gravity) were generated by shortening contraction (concentric). These kinematic and muscular patterns have been shown to reflect optimal solutions that take advantage gravity torque to minimize muscle effort during vertical movements of the upper limb (Gaveau et al. 2014, 2016, 2021; Berret et al. 2008; Crevecoeur et al. 2009). More interestingly, we found that the overall corticospinal output was reduced during lengthening compared to shortening contractions. This neural organization was produced by supra-spinal intracortical inhibition mechanisms as motoneuronal responsiveness remained unchanged between lengthening and shortening contractions. These data highlight a specific involvement of intracortical circuits during the neuromuscular control of vertical arm movements.

### Neural mechanisms implicated in gravity-oriented arm movements

Neurophysiological data from Experiment A showed a significant reduction of MEP amplitude in BB during downward (lengthening contraction) as compared to upward (shortening contraction) movements. This result copes with those of studies using force-generation tasks that systematically show downregulation of corticospinal excitability during lengthening contractions – whether the task involved maximal or submaximal contractions of upper or lower limb muscles (Abbruzzese et al., 1994; Sekiguchi et al., 2001; Duclay et al., 2011 and 2014; Gruber et al., 2009; Opie and al. 2016). It is important to note that the drop in corticospinal output could not be due to muscular mechanisms *per se* because both the coactivation and the RMS/M_max_ ratio in BB remained comparable between the two contraction modalities. Instead, it must have relied on supraspinal and/or spinal mechanisms.

To assess changes at the spinal level, we measured the responses to CMS and found that normalized CMEP amplitudes did not differ between downward and upward movements, thus indicating that motoneuron responsiveness to synaptic inputs remained similar across lengthening and shortening contractions. Consequently, the drop in global corticospinal excitability without any modulation of CMEP indicates the involvement of cortical mechanisms. The CMS is the most direct method to test motoneuron responsiveness to synaptic input in conscious humans (Martin et al., 2008). The CMEP has a large monosynaptic component in the upper limb (Petersen et al., 2002) and the descending tracts are not subject to presynaptic inhibitory mechanisms (Nielsen and Petersen,1994; Jackson et al., 2006).

Our results reveal that the neural commands responsible for opposite movement orientations, in the vertical plane, are tuned by intracortical mechanisms. Indeed, in Experiment B, both the SP and SICI indicated an increase in intracortical inhibition during lengthening contractions. The SP was longer in this contraction mode, even when MEP amplitudes were controlled – eliminating potential biases regarding the SP dependency to MEP size (Orth & Rothwell, 2004). SICI was also higher during lengthening contractions. Crucially, our paired-pulse protocol also allowed to assess SICI without confounding effects about the size of test MEPs, as corticospinal excitability was matched between conditions by adjusting TMS intensity. Without this procedure, the systematic variations in test MEP amplitude due to the muscle contraction mode (Duchateau and Enoka, 2008) make SICI protocols unreliable (Lackmy and Marchand-Pauvert, 2010; Opie and Semmler, 2014), which confounds the interpretation of the data, as in our Experiment A. Moreover, as our conditioning CMEP protocol showed that conditioning (subthreshold) MEPs did not reach the spinal level, SICI data can be safely considered as an accurate assessment of intracortical inhibitory circuits. SP and SICI are thought to reflect the contribution of GABA_B_-mediated and GABA_A_-mediated inhibition, respectively (Kimiskidis et al., 2006; Di Lazzaro et al., 1998). Interestingly, modulations of GABAergic inhibitory neurotransmission have been reported in the control of finger tracking tasks implying anisometric contractions, with a specific augmentation of both SP and SICI during lengthening contractions of intrinsic hand muscles (Sekiguchi et al., 2003; Opie and Semmler, 2016). Our findings provide strong evidence for a contraction-dependent modulation of intracortical inhibition in the control of vertical arm movements, where the neural command is specifically downregulated at the cortical level during lengthening contractions.

### Task dependent neural control of vertical arm movements

The characteristics of the motor task (e.g. type of muscle contraction, range, and speed of motion, level of external and/or internal forces) is a central issue when trying to identify the neural mechanisms of movement control. In many experiments, these mechanisms were evaluated through isokinetic actions where muscular contractions are induced by resisting a torque imposed by an ergometer, or by displacing a load to match a trajectory (Duchateau & Enoka, 2008; 2016). In such tasks, lengthening contractions specifically reveal spinal inhibitions (Nordlund et al., 2002; Sekiguchi et al., 2003; Gruber et al., 2009; Duclay et al., 2011 and 2014), which led to the interpretation that a relative increase in cortical excitability would compensate for it. Duclay et al. (2011, 2014) have indeed shown that the SP in the ongoing EMG recorded after a MEP was shorter during lengthening compared with shortening contractions of ankle flexors, suggesting a specific release of cortical inhibition. As shown by Inghilleri et al. (1993), silent periods longer than 100 ms are indeed mainly produced by cortical mechanisms. Also, Gruber and colleagues (2009) hypothesized a greater cortical excitability in lengthening contractions, because of larger MEP-to-CMEP ratios during lengthening compared with shortening contractions in the elbow flexors. Taken together, these studies suggest that the overall drop in corticospinal output during such lengthening contractions arises from an interaction between spinal and cortical mechanisms, where an extra excitatory drive from the motor cortex may be downregulated by spinal inhibitory mechanisms (see Duchateau & Enoka, 2016 for a review). In this context, the spinal modulations may highlight peripheral control loops that are required for the online regulation of the motor output, e.g. force, position (Scott, 2012; Nielsen, 2016). Low reflex gain favors the stability of muscle activation (Llewellyn et al. 1990), notably by mitigating the augmentation of Ia afferent activity due to proprioceptive inputs (Nielsen and Kagamihara 1993).

Our study highlights a distinct control strategy for natural, unconstrained vertical movements. It is worth mentioning that the modulation of intracortical circuits in the control of shortening and lengthening contractions during such a motor task may be related to the integration of gravity force, supporting predictive mechanisms of motor control (Franklin and Wolpert, 2011; Berret and Jean 2020). In fact, many behavioral and computational studies have proposed that the kinematic and EMG features of upward and downward movements are set at the motor planning stage (Papaxanthis et al., 2003; Gentili et al., 2007; Gaveau et al., 2014, 2016, 2019, 2021). A striking example of this, is the persistence of directional asymmetries during early adaptation to a microgravity environment (Papaxanthis at al., 2005; Gaveau et al., 2016). Although the load force is absent, it takes several trials before directional asymmetries progressively disappear, thereby converging towards newly optimal motor patterns (Gaveau et al. 2016). This result has been taken as the demonstration that a gravity internal model is recalibrated, and that deterministic optimal motor control is set predictively. Interestingly, these findings further support those of previous studies showing that the gravity internal model is supposed to be stored at the supra-spinal level, involving computations of the cerebellum, the anterior thalamus, the vestibular nuclei and the vestibular cortex (Angelaki et al., 2004; Laurens et al. 2013, 2016; Indovina et al. 2005; Rousseau et al., 2016).

## Conclusion

The neural policy we identified in the present study, i.e. a consistent contraction-dependent modulation of the intracortical inhibition while spinal excitability remained unchanged, well support that the hypothesis that gravity-related optimal motor commands are processed upstream to the muscle and spinal levels, within brain intracortical circuits.

## METHODS

### Participants

Sixteen healthy right-handed adults participated in this study (13 males and 3 females, aged between 23 and 48, mean age= 33.11 ± 6.34 years old). All were volunteers without any neurological or muscular disorders. Informed consents were signed, the study was approved by the Regional Ethics Committee of Bourgogne and performed in accordance with the Declaration of Helsinki.

### Study design

Two experiments were completed to assess corticospinal and spinal excitability modulation during single-joint (elbow anatomical angle) vertical movements in the sagittal plane. Ten participants were involved in experiment A and nine in experiment B (3 participants took part in both experiments with at least one-week interval between them). The motor task was strictly identical in both experiments; only the stimulation parameters for the neurophysiological measurements differed. Durations were ~2h30 and ~1h30 for experiments A and B, respectively. The experiments were carried out during the afternoon (between 1:00 p.m. and 5:00 p.m.) to control potential circadian effects (Gueugneau et al. 2010). On the day of the experiments, participants were required not to practice any sport or physical activity that could have altered their neuromuscular system.

### Motor tasks

Participants performed single-joint, visually guided pointing movements in the parasagittal plane with their right forearm (rotation around the elbow; see Figure 1, left panels). We chose one degree of freedom (DOF) movements to isolate the mechanical effects of gravity (Gentili et al. 2007; Le Seac’h and McIntyre 2007; Gaveau et al. 2014). Precisely, during single-joint vertical forearm movements, inertia (i.e., the distribution of the forearm mass around the elbow joint in a body-fixed coordinate system) remains constant, and inertial torque is only related to joint acceleration. Conversely, gravity torque significantly changes according to the movement direction. Note that during single-joint movements, interaction torque may also influence motion dynamics. For example, during the motion of the elbow joint, inertial interaction torques may arise at the shoulder and wrist joints because of elbow acceleration and deceleration. We confirmed that joint motion was restricted to the elbow joint only (see *Data analysis* below). At the muscular level, the task allowed us to focus on the Biceps Brachii (BB), which was activated in the two contraction modalities, according to the direction of the movement; i.e., lengthening (downward movement) and shortening contraction (upward movement). Fig 1 (right panels) qualitatively illustrates activation patterns from the BB and TB (triceps brachii) muscles. It is noticeable that upward and downward forearm movements are realized by the activation of the BB only.

Participants sat in a comfortable chair with their trunks vertically aligned and supported by the back of the chair. Their right upper-arm was in the vertical plane during the whole experiment. Three targets (1 cm diameter plastic spheres) were centered on the participants’ right elbow and positioned at a distance slightly superior to their forearm segment’s length. The initial target (IT) was horizontally aligned with the elbow. The other two targets were placed at an angle of 45° upward (UT) and −45° downward (DT), taking as reference the elbow-IT horizontal line (0°). We considered three different actions: one static and two dynamics. During the static action, the participants pointed towards the IT; the upper arm-forearm angle was 90°. This action involved an isometric contraction of elbow flexor muscles against gravity (i.e., the muscle was contracted without changing its length; muscle torque was equal to gravity torque) and was chosen to adjust parameters for magnetic and electrical stimulations (see details below). The dynamic actions comprised downward (with gravity) and upward (against gravity) movements. For the downward movements, the participants initially pointed to the UT during 2-3 seconds (the elbow was flexed at 45° and the semi-pronated hand was aligned with the forearm) before performing a movement to the DT. Note that this movement involved an eccentric contraction of elbow flexor muscles (i.e., the muscle is contracted and lengthened; its torque was inferior to gravity torque allowing downward motion of the forearm). For the upward movements, the participants initially pointed to the DT during 2-3 seconds (the elbow was flexed at −45° and the semi-pronated hand was aligned with the forearm) before performing a movement to the UT. Note that this movement involved a concentric contraction of elbow flexor muscles (i.e., the muscle is contracted and shortened; its torque was superior to gravity torque allowing upward motion of the forearm). For both movement directions, participants were informed that final accuracy was not the primary goal of the task. We trained participants (~ 10 practiced trials) to carried out upward and downward movements of ~0.5 s. An electronic metronome was used during the experiment to help them maintain it. We chose this speed because previous studies from our laboratory showed that at this velocity, participants accomplished movements at a natural and comfortable speed (Gaveau et al. 2011, 2014).

### Kinematics Recording

Kinematics was recorded using an optoelectronic device (VICON, Oxford, UK). Three cameras (100 Hz sampling frequency) were used to record the displacements of five reflective markers (1 cm in diameter) placed on the shoulder (acromion), elbow (lateral epicondyle), wrist (in the middle of the wrist joint between the cubitus and radius styloid processes), hand (first metacarpophalangeal joint), and the nail of the index fingertip.

### Electromyography

We recorded the electromyographic signals (EMG) from both the BB and the TB of the right arm. Two silver chloride (AgCl) surface electrodes (8 mm diameter; inter-distance 2cm) were positioned on both muscles after shaving and cleaning the skin. The electrodes were centered over the muscle bellies (lateral head of the TB; and medial part of the BB, ~2 cm above the elbow tendon). A common reference electrode was placed over the medial epicondyle of the left arm. EMG signals were amplified with a bandwidth frequency ranging from 15 to 5 kHz (gain: 1000), then digitized online (sampling frequency: 2 kHz), and stored into a personal computer for offline analysis using the MP150 acquisition system (BIOPAC Systems Inc., Santa Barbara, CA, USA).

### Neuromuscular stimulation methods

We used different stimulation methods in experiments A and B to evaluate distinct neurophysiological processes. In experiment A, we evaluated corticospinal excitability using single-pulse TMS, short intracortical inhibition (SICI) by paired-pulse TMS, and spinal motor neuron excitability through cervicomedullary stimulation (CMS). In experiment B, corticospinal excitability and SICI were evaluated by single-pulse and paired-pulse TMS, respectively, but with specific stimulation parameters allowing to match motor evoked potentials amplitude (MEP) across conditions (see details below). In both experiments, BB muscle excitability was assessed through Mmax recordings, which allowed to normalize the other electrophysiological variables.

#### Experiment A

##### Mmax recordings - brachial plexus stimulation

Single electrical stimuli were delivered to the brachial plexus to evoke M_max_ in BB (pulse duration 1 ms; provided by a Digitimer stimulator - model DS7; Hertfordshire, UK). The cathode was placed in the supraclavicular fossa and the anode on the acromion. To induce M_max_, the intensity was progressively increased (0.5-mA steps) from the perceptual sensory threshold to M_max_. Then, this intensity was further increased by ~20% to ensure supramaximal stimulation. Once determined at rest, the M_max_ intensity was then used to record M-waves during vertical forearm movements. Four M_max_ were recorded for each direction (upwards and downwards).

##### MEP and SICI recordings - transcranial magnetic stimulation

TMS was delivered to the optimal scalp position over the left motor cortex (M1) to activate the right BB. MEP was elicited by magnetic stimuli provided from a Bistim module combining two Magstim 200 stimulators (Magstim Company Ltd., Whitland, UK) through a figure-eight coil (loop diameter, 8 cm) with a monophasic current waveform. The coil was held tangentially to the scalp with the handle pointing backward and 45° away from the midline to activate the corticospinal system preferentially trans-synaptically via horizontal corticocortical connections (Di Lazzaro et al. 2004). The cortical representation of the BB was initially assessed with the stimulator intensity at 70% of its maximum stimulator output (MSO; 2.2 T). The optimal location was searched by slightly moving the coil over the M1 area until MEP of maximal amplitude and lowest threshold were recorded in the right BB. The optimal coil location was then marked on the participants’ scalp. As resting motor threshold (RMT) in proximal muscles could be hard to find in some participants, we used active motor threshold (AMT) to set TMS intensity. AMT was defined as the minimum intensity to produce a MEP amplitude of ≥ 300 μV in three out of five trials during weak muscle contractions (Rothwell et al., 1999). To do so, TMS was given while the participants pointed to the IT, thus producing a slight isometric contraction of the right BB. We confirmed in pre-experiment that the force developed during this postural configuration corresponded to ~3-5% of maximal voluntary contraction. Stimulation intensity was then set at 120% of the AMT to record single MEP during vertical arm movements (TMS intensity ranged from 51% to 78% of MSO; mean 62.1 ± 7.2%). The SICI protocol used a subthreshold conditioning pulse set at 80% of AMT given before the test stimulus (120% AMT) with an interstimulus interval (ISI) of 3 ms (Kujirai et al., 1993). Importantly, we confirmed during isometric contractions (static position) that the paired-pulse method effectively reduced MEP amplitude. Indeed, the mean MEP amplitude of the conditioned responses (0.33 ± 0.05 mV) was significantly lower (P < 0.05, t(9) = 10.99) than the unconditioned responses (0.61 ± 0.09 mV). Corticospinal excitability and SICI were evaluated by recording unconditioned and conditioned MEP during vertical forearm movements. Ten unconditioned MEP and 10 conditioned MEP were recorded for each direction (upwards and downwards).

##### Cervicomedullary stimulation (CMS)

CMS was used to directly measure spinal motoneuron excitability by eliciting a single volley in descending axons at the pyramidal decussation level (Taylor 2006). It is known that CMEP is challenging to record in some participants, due to the discomfort induced by the stimulation (Taylor et al. 2002). Here, we discarded two participants as they presented noise in the EMG signal (i.e., slight visually detectable EMG activities at rest) due to apprehension and general discomfort. CMS was given by placing two AgCl (2 cm diameter) electrodes over the mastoid processes on both sides with the cathode placed at left. We used a DS7AH current stimulator (Digitimer, Hertfordshire, UK) to elicit 200 μs-width pulses. The RMT was determined individually as the minimal intensity to evoke peak-to-peak CMEP amplitude of 50 μV in the resting BB. The intensity was then set to 120% of the participant’s RMT (mean intensity: 195.5 ± 65 mA; range: 110–270 mA) to record CMEP during vertical forearm movements. Four CMEP were recorded for each direction (upwards and downwards).

We also provided a complementary measure of spinal excitability by conditioning CMS with a TMS pulse over M1. When provided in close temporal proximity, CMS and TMS interact at the spinal level and give rise to a complex motor response in a target muscle. Precisely, when a supra-threshold TMS pulse precedes CMS by 5 ms, responses to paired stimuli are larger than the responses to individual stimuli (Taylor et al., 2002). This indicates that the signal from TMS interacts with the spinal motoneuron pool before the arrival of the volley from CMS and thus induces a facilitatory motor response. More, we have previously shown that a sub-threshold M1 magnetic stimulation could modulate spinal excitability during a force-generation task (Grospretre et al., 2014). These data are crucial when one considers SICI set-ups, as the conditioning TMS pulse may influence spinal excitability, biasing thus the interpretations about the underlying neural mechanisms. To tackle this issue, we recorded CMEP during a paired-pulse TMS-CMS protocol (Taylor et al., 2002), where a sub-threshold TMS pulse (80% AMT, as in our SICI protocol) was given 5 ms before CMS. So, supposing that paired stimuli and single CMS induce motor responses of similar size, it could be concluded that the sub-threshold TMS pulse does not affect spinal excitability and that our SICI protocol does properly assess intracortical mechanisms. TMS coil manipulation and CMS arrangement were the same as detailed above. Four conditioned CMEP were recorded for each forearm movement (upwards and downwards).

#### Experiment B

Experiment B was specifically designed to assess intracortical inhibition while controlling corticospinal excitability across the experimental conditions, where test MEPs in the SICI protocol were carefully matched between shortening and lengthening contractions.

##### Mmax recordings - brachial plexus stimulation

The procedure was identical to experiment A, with M_max_ being recorded during vertical arm movements (stimulation intensities ranged from 3.2 to 19.4 mA). Four M_max_ have been recorded for each upward and downward arm movement.

##### MEP and SICI recordings - transcranial magnetic stimulation

Coil location and AMT were similar as in experiment A. MEP and SICI were first recorded during isometric contractions (as in experiment A), with stimulation intensities at 120% and 80% of AMT for test and conditioning pulses, respectively. At the group level, unconditioned and conditioned MEP amplitudes in isometric condition were 0.51 ± 0.07 mV and 0.28 ± 0.04 mV, respectively (P < 0.05, t(8) = 11.7). Then, TMS intensity was adjusted to produce unconditioned (test) MEP of similar amplitudes (~0.5 mV; as in isometric condition) for upward and downward movements. Stimulation intensity thus had to be slightly lowered for both upward (shortening contractions) and downward movements (lengthening contractions) compared to isometric contractions; i.e., 62.3 ± 7.2% of MSO in isometric contraction vs. 58.1 ± 5.2% and 61.4 ± 5.2% for shortening and lengthening contractions, respectively. The conditioning TMS pulse remained at 80% of AMT during the whole experiment. Thanks to this procedure, unconditioned and conditioned MEP were recorded during upward and downward movements, while the corticospinal output was matched across conditions. Ten unconditioned MEP and 10 conditioned MEP were recorded for each arm movement (upward and downward).

### Experimental conditions and recording procedure

In both experiments, before performing the trials including neurophysiological stimulations, the participants performed 24 trials (12 upwards, 12 downwards) without any stimulation. This procedure allowed us to evaluate the arm kinematics and EMG signals without the stimulation artifact. Then, the neurophysiological measures were carried out in a block design. The recording order of the variables was counterbalanced across participants. For instance, for a given participant in experiment A, the order of data collection could be: M_max_, unconditioned MEP, conditioned MEP, CMEP, and conditioned CMEP for upwards and downwards arm movements; and then changed for the next participant. Experiment B included the same variables except for CMEP. Sixty-four trials with neurophysiological measures were performed in experiment A [(4 M_max_, 10 unconditioned MEP, 10 conditioned MEP, 4 CMEP, 4 conditioned CMEP) x 2 (upwards, downwards movements)], and 48 in experiment B [(4 M_max_, 10 unconditioned MEP, 10 conditioned MEP) x 2 (upwards, downwards movements)]. Among these trials, one trial free from any stimulation for 4 trials with neurophysiological measures was added. These trials were inserted randomly to prevent potential habituation and anticipation effects due to electrical/magnetic stimulations. A ten min break was included approximately in the middle of each experimental session to prevent any fatigue effect.

Stimulations for both upwards and downwards movements were given when the arm and forearm reached a 90° angle. This guaranteed that arm configuration and muscle length were similar whatever the movement direction. The timing of stimulations was precisely triggered using a two-axis electronic goniometer (Biometrics Ltd, SG110) attached to the participants’ right elbow (epicondyle). Arm/forearm angular displacements in the sagittal plane were recorded and monitored online (signal sampling frequency: 2 kHz). The goniometer calibration was undertaken before each experimental session and cautiously checked throughout the experiments. The triggering delay was ~1 ms (BIOPAC Systems Inc., Santa Barbara, CA, USA).

### Data analysis

#### Movement Kinematics

This analysis concerned trials free from any stimulation. Data processing was processed using custom MATLAB programs (MathWorks, Natick, MA). Kinematic signals were low-pass filtered (5-Hz cutoff frequency) using a digital fifth-order Butterworth filter (zero-phase distortion, “butter” and “filtfilt” MATLAB functions, Mathworks). Three-dimensional velocity signals were inspected to ensure that they were single-peaked. Angular displacements were inspected to verify that participants performed one DOF movements with the elbow joint. Joint movements were discarded from further analysis when they showed multiple local maxima and/or a rotation (≥3°) to other than the elbow joint (~2% of all trials). Movement onset and offset were defined with a threshold of 10% of the maximal angular velocity (Gaveau el al. 2011).

We calculated the following kinematic parameters from the index fingertip: movement amplitude and duration, mean and peak velocity, time to peak velocity, peak acceleration, and time to peak acceleration. From these variables, *invariant* parameters were computed: i) relative time to peak velocity (rtPV = time to peak velocity / movement duration) and ii) relative time to peak acceleration (rtPA = time to peak acceleration / movement duration). These parameters are termed *invariant* because they could remain constant across experimental conditions and are theoretically independent of movement direction, speed, and amplitude (Atkeson and Hollerbach 1985; Gaveau et al., 2016). Here, they were used as behavioural markers to examine whether participants produce physiological patterns during vertical movements, as previously revealed by specific kinematics for upward and downward movements (Papaxanthis et al., 2005; Gaveau et al., 2016). Besides, to make qualitative comparisons between directions, we normalized the velocity profiles in time (cubic spline function; Math-Works) and amplitude (velocity time series divided by maximal velocity). Normalization guarantees that velocity profiles are independent of joint amplitude, time, and maximal velocity.

#### EMG activity

The procedure was identical for both experiments. The EMG signal was used to i) characterize the EMG activity of BB and TB during upward and downward movements and ii) quantify the electrophysiological responses to neuromuscular stimulations; i.e. M_max_, MEP, silent period (SP), and CMEP.

BB and TB EMG activity patterns were characterized by computing the root mean square (RMS) for every 50 ms period of the EMG signal. BB RMS values of the EMG signal over a 50 ms period before the stimulation were normalized to the mean amplitude of the M_max_ (RMS/M_max_) for both movement directions. This normalization procedure was used to control contraction-related changes at the muscle level (Duclay et al., 2011). Muscular coactivation was expressed as a ratio of antagonist to agonist muscle activity (Hu et al., 2007; Rao et al., 2009), by normalizing TB RMS value to BB RMS value within the 50 ms preceding the stimulation. For each experiment, a mean value of coactivation was computed over all trials for each movement condition and participant.

#### Evoked potentials

For each electrophysiological measure, we considered the peak-to-peak amplitude of the EMG response. M_max_, MEP, and CMEP amplitudes from BB were averaged for each movement, condition, and participant. Considering MEP, trials in which MEP amplitude was more prominent than two SDs were considered as outliers (see Methods in Gueugneau et al. 2015). On average 0.9 (SD: 1.2) out of 10 MEP were excluded. SICI was expressed as a difference between the conditioned and unconditioned test MEP and quantified in percentage: SICI = [(MEP_conditioned_ - MEP_unconditioned_ / MEP_unconditioned_) × 100]. Thus, high and low negative values indicate strong and weak SICI, respectively, while positive values would indicate a facilitation. The same formula was then used to quantify the conditioned CMEP amplitude in our paired-pulse TMS-CMS trials (Experiment A). Changes in intracortical inhibition were also evaluated by measuring SP in the ongoing EMG following TMS. The SP duration was taken as the time interval from the stimulus artifact to the return of continuous EMG (Damron et al., 2008). The end of the SP was determined when the corresponding rectified EMG activity reached a value within two SD of the rectified mean EMG signal recorded during 1 s when the participant was at rest. Because the duration of the SP can be influenced by the size of the MEP (Orth and Rothwell, 2004), we also computed the relation between the two parameters (SP/MEP ratios) for both shortening and lengthening contractions. Finally, for both experiments and contraction types, MEP and CMEP amplitudes of the BB were normalized to the corresponding Mmax amplitude obtained in the same condition, to reduce inter-subject variability and to reliably evaluate contraction-dependent changes in the corticospinal network (Lackmy and Marchand-Pauvert 2010; Palmieri et al., 2004).

### Statistics

All data are presented as means ± standard error (SE). The normality of the data was confirmed using the Shapiro-Wilks W test (*P* > 0.05). First, our study aimed to confirm the kinematic differences between upward and downward arm movements previously described in the literature. For that reason (unilateral hypothesis), all kinematics variables were submitted to one-tailed paired *t-tests* (upward vs. downward). Direction-dependent differences in electrophysiological variables were the second step in our analyses. All electrophysiological variables were submitted to two-tailed paired *t-tests* (upward vs. downward). Multi-factor repeated measures analysis of variance (ANOVA) was used when necessary. The effect size was evaluated by calculating the Cohen’s *d*, and partial eta-squared (η_p_^2^) for *t-*tests comparison and ANOVA, respectively. Significance was set at *P* < 0.05.

## ACKNOWLEDGEMENTS

This work was supported by the French “Investissements d’Avenir” program, project ISITE-BFC (contract ANR-15-IDEX-0003). We would like to thank Yves Ballay for his help with data acquisition and technical support.

## COMPETING INTERESTS

The authors declare that they have no financial competing interests.

## Notes

### Competing Interest Statement

The authors have declared no competing interest.

